## Supplementary Materials for "The DNAPKcs long-range C-NHEJ complex is required for blunt DNA end joining when XLF is weakened"

### Supplemental Table S1. Oligonucleotide list.

| **Name** | **Purpose** | **Sequence (5' → 3', all sgRNA sequences have the initial G nucleotide, regardless of whether it is part of the targeted sequence)** |
|---|---|---|
| 7a | sgRNA | GACCACCCTGACCTACGGCTA |
| 7b | sgRNA | GGCTGAAGCACTGCACGAAT |
| 7a+1 | sgRNA | GTAGATATCTTCCTTAGCCGT |
| GAPDH | sgRNA | GTATAGAAACCGGGGGCGCGG |
| CD4 | sgRNA | GGCGTATCTGTGTGAGGACT |
| DSB-H | sgRNA | GACGCCCCAGCACTCGTCCGA |
| DSB-G | sgRNA | GAGCACTGCACGCCGTAGGTC |
| DSB-L | sgRNA | GCTCTTCGCTATTACGCCAGC |
| mXlfsg1 | sgRNA | GCAAAACAGCAGATCCAAGCA |
| mXlfsg2 | sgRNA | GTCAAGCAGCACCTCCCCTCG |
| PRKDCsg1 | sgRNA | GACTAAAGGCAATTCGTCCTC |
| PRKDCsg2 | sgRNA | GAGACACGTAGTTGTCCAGA |
| XLFsg1 | sgRNA | GTTGGTTTCAGATCTTCAAC |
| LMNA | sgRNA | GCCATGGAGACCCCGTCCCAG |
| ILL-GAPDH | primer | ACACTCTTTCCCTACACGACGCTCTTCCGATCTACGTAGCTCA GGCCTCAAGA |
| ILL-CD4 | primer | GACTGGAGTTCAGACGTGTGCTCTTCCGATCTACAGTTCCAGT GGGAAATCG |
| *TIDE analysis primers for GFPd2-SV* | | |
| LACZ4pcrUP1 | primer | AACTTCAAGCTTGGCACTGG |
| LACZ4pcrDN1 | primer | GACAGTATCGGCCTCAGGAA |
| HYG2pcrUP1 | primer | GACGGCAATTTCGATGATG |
| HYG2pcrDN1 | primer | CGAAGCCCAACCTTTCATAG |
| GFP1pcrUP1 | primer | ACGTAAACGGCCACAAGTTC |
| GFP1pcrDN1 | primer | AAGTCGTGCTGCTTCATGTG |
| *qPCR analysis primers for GFPd2-SV* | | |
| A site UP | primer | CGTACTACGAGATTTCGATTCCA |
| A site DN | primer | TTATAAGCTGCAATAAACAAGTTGG |
| B site UP | primer | TGTCTTGTGCCCAGGAGAG |
| B site DN | primer | CAACAGATGGCTGGCAACTA |
| C site UP | primer | TCCCTTTAGGGTTCCGATTT |
| C site DN | primer | GTTTTCCCAGTCACGACGTT |
| Actin UP | primer | ACTGGGACGACATGGAGAAG |
| Actin DN | primer | AGGAAGGAAGGCTGGAAGAG |

**Supplemental Figure S1. DNAPKcs analysis in U2OS cells. (a)** DNAPKcs is less important for No Indel EJ vs. XLF and XRCC4 in U2OS cells. n=6. Statistics with unpaired *t*-test using Holm-Sidak correction. Immunoblots show levels of DNAPKcs and XRCC4. **(b)** M3814 causes a dose dependent decrease for No Indel EJ in U2OS cells. n=6. Statistics as in (a). **(c)** Immunoblots show levels of XLF-WT and XLF-K160D in U2OS cells. ****$P$<0.0001. Error bars = SD.

**Supplemental Figure S2. GAPDH-CD4 insertions are consistent with Cas9 staggered DSBs causing 5' overhangs; and EJ7+1-GFP validation. (a)** Shown are predicted blunt EJ outcomes for the GAPDH-CD4 rearrangement following paired blunt DSBs (No Indel EJ), a staggered 1 nt 5' overhang DSB at GAPDH that is filled in (+1 C nucleotide insertion), and a staggered 2 nt overhang DSB at GAPDH that is filled (+2 CG nucleotide insertion). Staggered cleavage at the *CD4* gene would not be expected to cause GAPDH-CD4 insertion mutations, because the staggered DSB would remove nucleotides on the 3' strand of the distal end, which when filled in, restore the predicted blunt DSB. (**b**) Shown are the frequency of insertion sizes for the control HEK293 cells treated with DMSO. n=3. **(c)** For the samples shown in (b) is the frequency of each insertion size with the sequence consistent with Cas9 staggered DSBs causing 5' overhangs that is followed by fill-in synthesis and blunt EJ. n=3. **(e)** Shown is the EJ7+1-GFP reporter (not to scale) and the Sanger sequence trace of GFP+ cells sorted from HEK293 parental cells using this assay. Error bars = SD.

**Supplemental Figure S3. HDR analysis in *XLF-KO/PRKDC-KO* cells, validation experiments for the GFPd2-SV reporter, and validation of the XLF-WT and K160D stable HEK293 cell lines. (a)** Shown is the frequency of HDR via the LMNA-HDR assay in *XLF-KO/PRKDC-KO* cells with EV or complementation vectors. n=6. **(b**) Validation of GFPd2-SV reporter in HEK293 cells. Shown is a diagram of the GFPd2-SV reporter (not to scale) with loci positions examined by qPCR (A, B, and, C, not to scale). GFP-negative (GFP-neg) cells were enriched by cell sorting following DSBs induced by the sgRNAs DSB-H, DSB-G, and DSB-L. Such GFP-neg cells were then examined for amplification of each locus shown (A, B, and C) relative to parental GFP+ cells, using Actin amplification to calculate $2^{-\Delta\Delta Ct}$. n=3 amplifications. Shown is TIDE analysis of samples following DSBs induced by sgRNAs DSB-H, DSB-G, and DSB-L, each using amplification with primers flanking the predicted DSB site. n=3. Error bars = SD. *P*<0.05, *******P*<0.01, ***********P*<0.0001, n.s. = not significant. One Way Anova with Tukey's post-test. **(c)** Validation of HEK293 stable cell lines using the EJ7-GFP reporter. n=6. Error bars = SD. *****P*<0.0001, unpaired *t*-test using Holm-Sidak correction. Immunoblots show levels of XLF-WT and XLF-K160D for these stable cell lines.

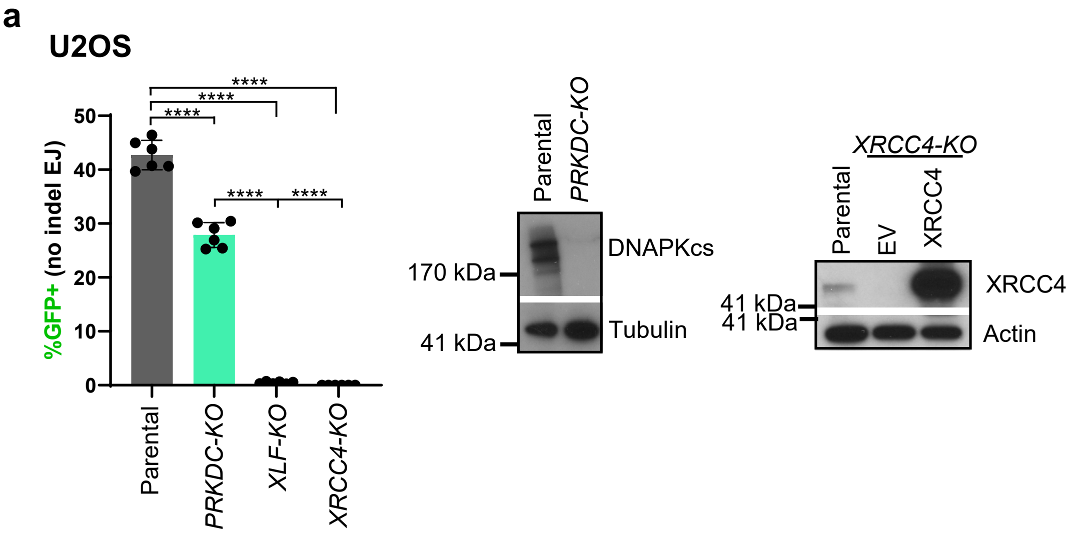
a
U2OS
%GFP+ (no indel EJ)
****
Parental
PRKDC-KO
XLF-KO
XRCC4-KO
DNAPKcs
Tubulin
170 kDa
41 kDa
XRCC4-KO
EV
XRCC4
Actin

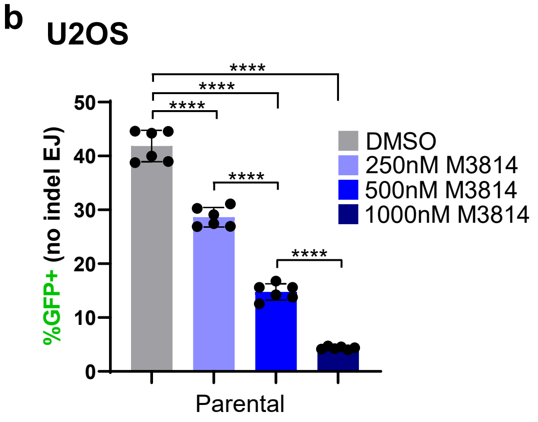
b
U2OS
%GFP+ (no indel EJ)
****
DMSO
250nM M3814
500nM M3814
1000nM M3814
Parental

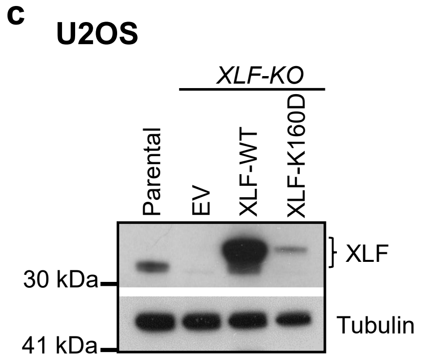
c
U2OS
XLF-KO
Parental
EV
XLF-WT
XLF-K160D
XLF
Tubulin
30 kDa
41 kDa

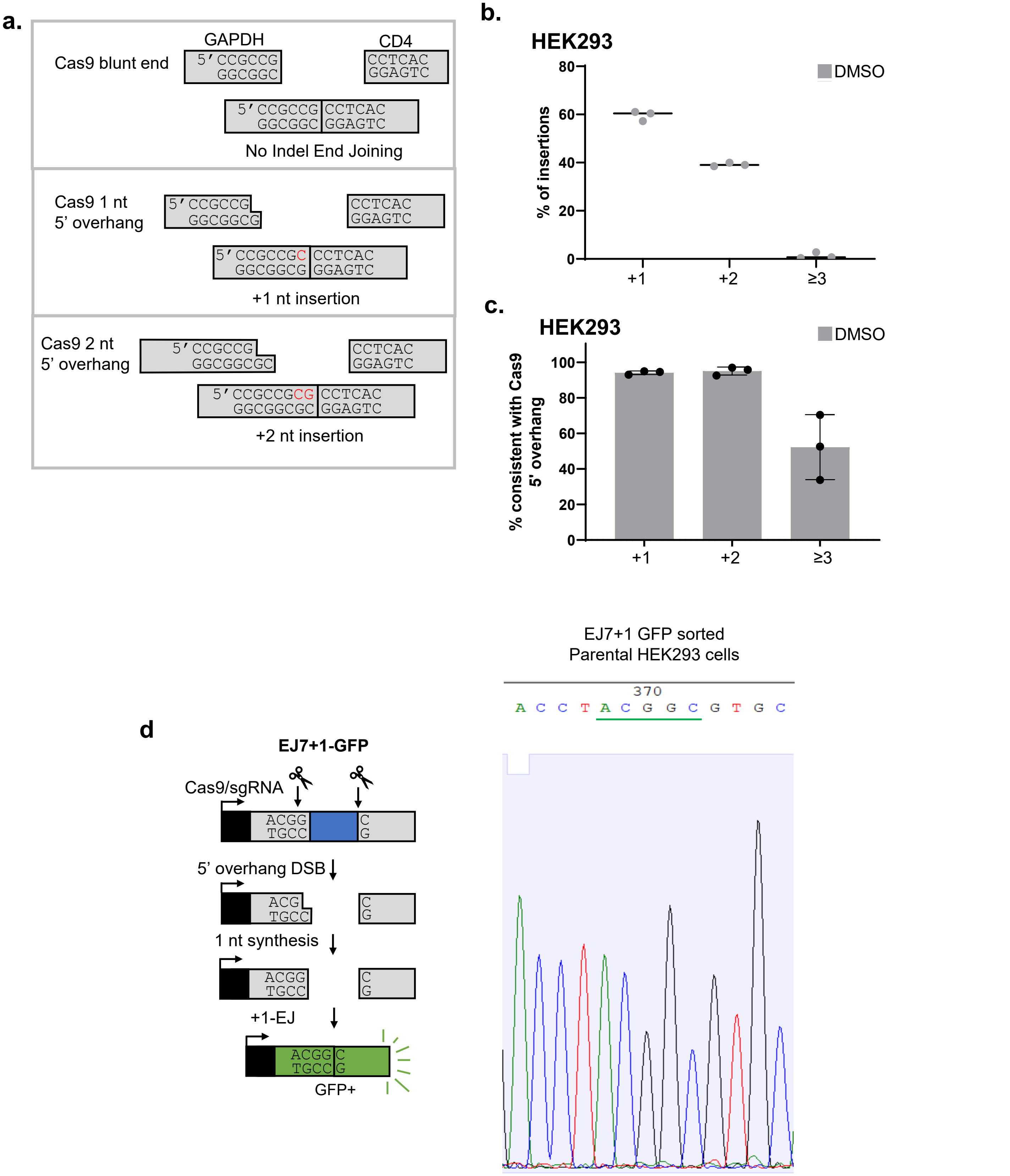
a.
Cas9 blunt end
GAPDH
5' CCGCCG
GGCGGC
CD4
CCTCAC
GGAGTC
5' CCGCCG CCTCAC
GGCGGC GGAGTC
No Indel End Joining
Cas9 1 nt
5' overhang
5' CCGCCG
GGCGGCG
CCTCAC
GGAGTC
5' CCGCCGC CCTCAC
GGCGGCG GGAGTC
+1 nt insertion
Cas9 2 nt
5' overhang
5' CCGCCG
GGCGGCGC
CCTCAC
GGAGTC
5' CCGCCGCG CCTCAC
GGCGGCGC GGAGTC
+2 nt insertion
b.
HEK293
DMSO
% of insertions
0
20
40
60
80
+1
+2
≥3
c.
HEK293
DMSO
% consistent with Cas9
5' overhang
0
20
40
60
80
100
+1
+2
≥3
d
EJ7+1-GFP
Cas9/sgRNA
ACGG
TGCC
C
G
5' overhang DSB
ACG
TGCC
C
G
1 nt synthesis
ACGG
TGCC
C
G
+1-EJ
ACGG C
TGCC G
GFP+
EJ7+1 GFP sorted
Parental HEK293 cells
370
A C C T A C G G C G T G C

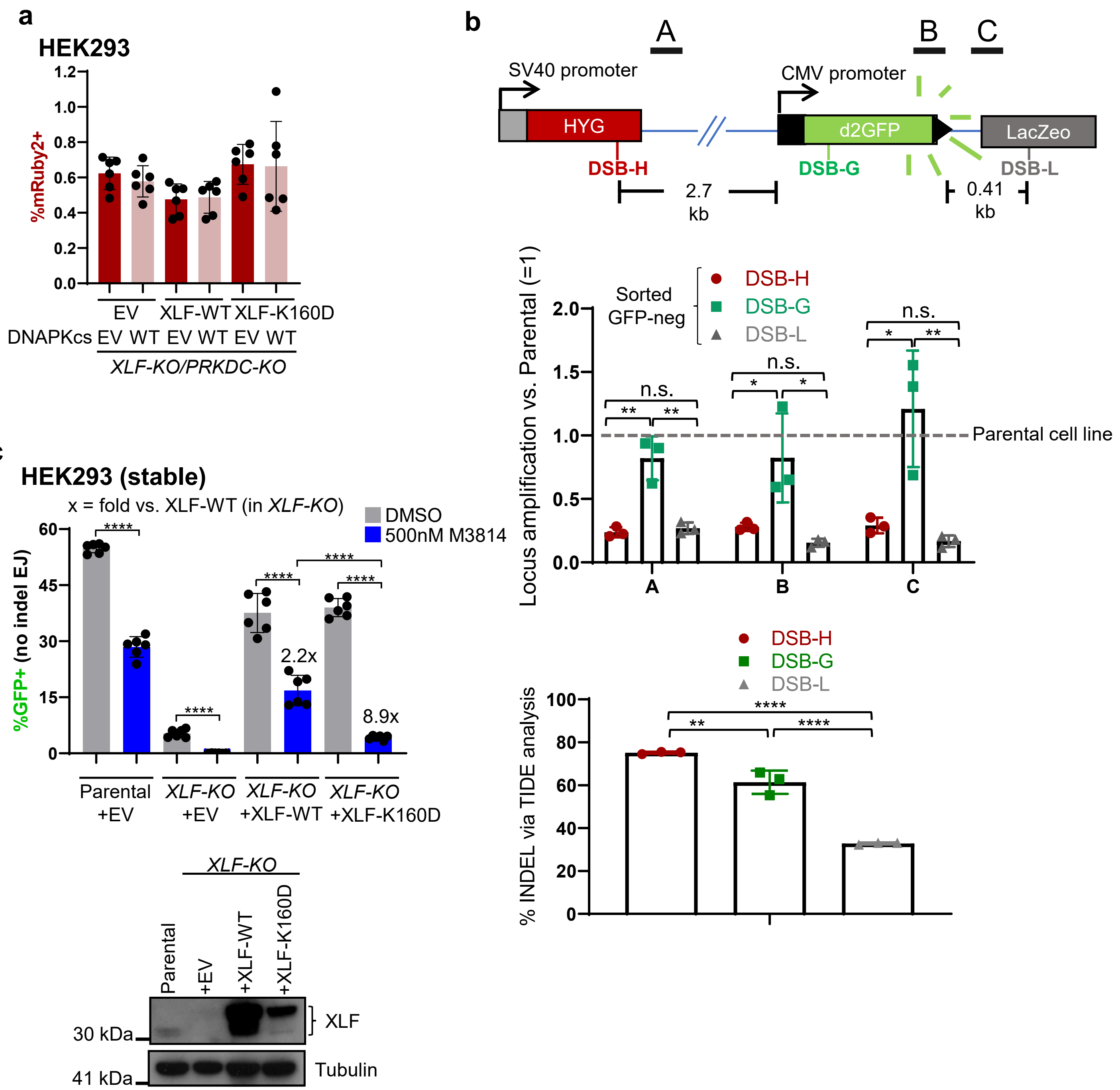
a
HEK293
%mRuby2+
EV
XLF-WT
XLF-K160D
DNAPKcs EV WT EV WT EV WT
XLF-KO/PRKDC-KO
b
A
B
C
SV40 promoter
CMV promoter
HYG
d2GFP
LacZeo
DSB-H
DSB-G
DSB-L
2.7 kb
0.41 kb
Sorted GFP-neg
Locus amplification vs. Parental (=1)
Parental cell line
n.s.
% INDEL via TIDE analysis
c
HEK293 (stable)
x = fold vs. XLF-WT (in XLF-KO)
DMSO
500nM M3814
%GFP+ (no indel EJ)
2.2x
8.9x
Parental +EV
XLF-KO +EV
XLF-KO +XLF-WT
XLF-KO +XLF-K160D
XLF-KO
Parental
+EV
+XLF-WT
+XLF-K160D
30 kDa
41 kDa
XLF
Tubulin
